# Gene expression associated with disease resistance and long-term growth in a reef-building coral

**DOI:** 10.1101/2020.08.12.248286

**Authors:** Emma R. Kelley, Robin S. Sleith, Mikhail V. Matz, Rachel M. Wright

**Affiliations:** Smith College, Department of Biological Sciences; University of Texas at Austin, Department of Integrative Biology

**Keywords:** coral disease, gene expression, *Montastraea cavernosa*, Flower Garden Banks National Marine Sanctuary

## Abstract

Rampant coral disease, exacerbated by climate change and other anthropogenic stressors, threatens reefs worldwide, especially in the Caribbean. Physically isolated yet genetically connected reefs such as Flower Garden Banks National Marine Sanctuary (FGBNMS) may serve as potential refugia for degraded Caribbean reefs. However, little is known about the mechanisms and trade-offs of pathogen resistance in reef-building corals. Here we measure pathogen resistance in *Montastraea cavernosa* from FGBNMS. We identified individual colonies that demonstrated resistance or susceptibility to *Vibrio spp*. in a controlled laboratory environment. Long-term growth patterns suggest no trade-off between disease resistance and calcification. Predictive (pre-exposure) gene expression highlights subtle differences between resistant and susceptible genets, encouraging future coral disease studies to investigate associations between resistance and replicative age and immune cell populations. Predictive gene expression associated with long-term growth underscores the role of cation transporters and extracellular matrix remodelers, contributing to the growing body of knowledge surrounding genes that influence calcification in reef-building corals. Together these results demonstrate that coral genets from isolated sanctuaries such as FGBNMS can withstand pathogen challenges and potentially aid restoration efforts in degraded reefs. Furthermore, gene expression signatures associated with resistance and long-term growth help inform strategic assessment of coral health parameters.

## INTRODUCTION

Infectious diseases cause mass coral mortality worldwide, especially in the Caribbean where Stony Coral Tissue Loss Disease (SCTLD) has massively reduced live coral cover (Precht et al. 2016; Walton, Hayes, and Gilliam 2018; Rippe et al. 2019). Several *Vibrio* species contribute to coral diseases, though the exact etiological agents for many outbreaks, including SCLTD, are uncharacterized (Weil, Smith, and Gil-Agudelo 2006; Chimetto Tonon et al. 2017). Heterogeneity in disease outcomes exists between and within coral species. For example, acroporid and pocilloporid coral species appear to be among the most vulnerable taxa (Hobbs et al. 2015) while massive corals, like *Porites*, resist bacterial challenge (Fuess et al. 2017). These species-level differences in disease resistance shape reef communities (Aronson, Precht, and Macintyre 1998; Williams and Miller 2012). Variation in disease susceptibility observed among members of a coral species (e.g., Wright et al. 2017, 2019; Libro and Vollmer 2016) may contribute to reef restoration if resistant genets can repopulate degraded reefs (van Oppen et al. 2017).

At 190 km off the Louisiana–Texas coastline, healthy corals in Flower Garden Banks National Marine Sanctuary (FGBNMS) produce larvae that can disperse throughout the Caribbean (Davies et al. 2017). This deep and isolated reef environment has maintained >50% coral cover with no documented disease outbreaks (Johnston et al. 2016). However, a highly localized mortality event occurred in late July 2016, affecting 5.6 ha (2.6% of the area) of the East Flower Garden Bank (FGB) while the West FGB remained unaffected (Johnston et al. 2019). Diverse invertebrates presented with advancing lesions of tissue loss that mimic an infectious disease. Studies found that localized hypoxia contributed to this disease-like mortality rather than a specific bacterial pathogen (Johnston et al. 2019; Kealoha et al. 2020). Given the importance of this sanctuary as a source population to help restore Caribbean reefs, it is critical to assess the ability of its coral inhabitants to withstand disease challenges.

Here we measure variation in susceptibility to a bacterial pathogen in the Great Star Coral *Montastraea cavernosa* from FGBNMS. Once considered among the most robust Caribbean species (Pinzón et al. 2014), *M. cavernosa* have experienced substantial mortality from SCTLD in recent years (Walton, Hayes, and Gilliam 2018). In addition to assessing susceptibility as the appearance of tissue loss upon challenge with *Vibrio spp*., we also measure long-term calcification to account for trade-offs between growth and resistance. This coral species has demonstrated stable calcification rate under heat stress (Manzello et al. 2015), but the impacts of disease on coral growth are unknown.

This study also characterizes predictive gene expression to identify molecular markers associated with long term calcification and resistance to bacterial pathogen invasion. Recent studies have made progress identifying allelic variation associated with coral thermal tolerance by sequencing hundreds of corals and often relying on reproductive crosses (Manzello et al. 2015; Jin et al. 2016; Fuller et al. 2020; Quigley, Bay, and van Oppen 2020). Here we rely on global gene expression, which can be used to associate gene expression with complex phenotypes in wild populations (Rose, Seneca, and Palumbi 2016). The genes the expression of which is associated with resistance to bacterial challenge and calcification over the subsequent year enhance our understanding of the molecular determinants of disease resistance and growth in coral.

## METHODS

### Coral Collection & Fragmentation

Coral fragments were obtained from the East and West Texas Flower Gardens Banks on 10 November 2014. SCUBA divers retrieved *M. cavernosa* fragments using hammers and chisels. Fifteen colonies were sampled from both the East and West banks for a total of 30 colonies. The larger sampled coral fragments were then divided with a wet saw into control and experimental series (n = 2–3 fragments per genet per series; mean±SD area = 4.8±1.8 cm^2^). The replicate fragments recovered in 15-gallon tanks of 32 ppt artificial seawater (ASW; Instant Ocean). Tanks were maintained at 23°C under 12,000K LED lights on a 12:12 hour light/dark cycle. Corals were fed Coral Frenzy every two days. After two days recovery, fragments were moved to individual experimental chambers containing 300 mL ASW under the same lighting and temperature conditions.

### Bacterial Challenge & Survival

Single isolates of *Vibrio coralliilyticus* or *V. shiloi* were incubated overnight in Difco Marine Broth-2216 (BD) along with a sterile broth control at 30°C with shaking (150 rpm). Overnight cultures were triple-washed in sterile ASW by centrifugation at 5000□*g* for ten minutes and resuspension in ASW. Corals were challenged with either 10^7^ CFU/mL of triple-washed *Vibrio* (treatment) or the same volume of triple-washed marine broth (control) on 28 November and subsequently every 24 hours for 17 days. We challenged the corals with *V. shiloi* for the first 10 days and *V. coralliilyticus* for the subsequent week of bacterial inoculations. The temperature was ramped to 29°C between the 5^th^ and 6^th^ days of inoculation. The fragments were photographed daily using a Nikon D5100 with a coral health card to monitor lesion development. Time-of-death was recorded as the day when tissue loss exceeded 50% of the surface area. After the 17^th^ day of bacterial challenge, surviving genets were placed in 15-gallon aquaria under control conditions.

### Growth Measurements & Analysis

The number of polyps were counted on each of the surviving fragments in February 2015 and again a year later, in March 2016. Surviving individuals were weighed in February 2015 and again in February 2016 following the buoyant weight protocol (Rose, Seneca, and Palumbi 2016; Davies and Spencer Davies 1989). Temperature and lighting conditions remained constant over the long-term monitoring period (25°C, 12:12 hour light/dark). Surface areas for each fragment were measured using ImageJ (Schneider, Rasband, and Eliceiri 2012). Skeletal weight and polyp growth were normalized to surface area. Statistical analyses were conducted in R version 3.6.1 (R Core 2019). The R package *MCMCglmm* (Hadfield 2010) was used to fit generalized linear mixed models for tissue and skeletal growth rates between phenotype (resistant vs. susceptible), treatment (control vs. *Vibrio*-challenged), and collection location (East vs. West Bank).

### Predictive Gene Expression

RNA was isolated from two replicate subsamples of each genet before bacterial challenge using the RNAqueous Total RNA Isolation Kit (Invitrogen). A total of 54 gene expression libraries prepared following the Tag-Seq protocol (Meyer, Aglyamova, and Matz 2011) were of high enough quality for Illumina HiSeq 2500 sequencing (SRA: PRJNA355872). Adapter sequences were trimmed and low quality reads (minimum quality score = 20; minimum percent bases above minimum quality score = 90%) were filtered using FASTX toolkit (Gordon, Hannon, and Others 2010). Reads were mapped to a holobiont reference consisting of the *M. cavernosa* genome (Fuller, Z., Yi, L., and Matz, M., 2020) and *Cladocopium goreaui* transcriptome (Davies et al. 2018) using Bowtie 2 (Langmead and Salzberg, 2012). Reads were converted to counts representing the number of independent observations of a transcript over all isoforms for each gene.

Isogroups (henceforth called “genes”) with a mean count less than three across all samples were removed from the analysis. Expression sample outliers were detected using *arrayQualityMetrics* (Kauffmann, Gentleman, and Huber 2008). Differentially expressed genes (DEGs) were identified using *DESeq2* (Love, Huber, and Anders 2014). Wald tests were performed to compare phenotype (resistant vs. susceptible) and collection location (EFGB vs. WFGB) using the model count ∼ pheno + bank. Wald tests were also performed to compare continuous growth phenotypes using the models count ∼ mean calcification rate and count ∼ mean polyp generation. We report Wald statistics (log fold change/standard error) to represent the magnitude of expression difference between groups or per unit change of continuous variables. False-discovery rate (FDR) *p*-values were adjusted using the Benjamini– Hochberg procedure (Benjamini and Hochberg 1995). Gene expression heatmaps were generated using *pheatmap* (Kolde 2012) and gene ontology enrichment was performed based on signed adjusted *p*-values using *GO-MWU* (Wright et al. 2015).

### Reference-based 2bRAD Genotyping

We prepared 64 genotyping libraries using the 2bRAD protocol (Wang et al. 2012) and sequenced the libraries on the Illumina HiSeq 2000 platform at UT Austin Genome Sequencing and Analysis Facility. We used FASTX toolkit to remove barcodes, deduplicate reads, and apply quality filters such that only reads in which 90% or more of the bases with a Phread score >= 20 were retained. These reads were mapped to the *M. cavernosa* genome (Fuller, Z., Yi, L., and Matz, M., 2020.) using bowtie2 (Langmead and Salzberg 2013). Genotyping was performed with ANGSD v0.930 (Korneliussen, Albrechtsen, and Nielsen 2014). Sites were filtered to retain loci with a mapping quality >= 20, base call quality >= 30, and minor allele frequency >= 0.05 that were sequenced in at least 20 individuals. These sites were used to calculate pairwise identity-by-state distances between individual samples. Distances <0.15 presumed to be clones based on similarity detected across genotyping replicates. Only one clone per sample retained for subsequent population genetic analysis. Library replicates removed as clones, as well as two additional pairs of clones. The VCFtools subprogram *weir-fst-pop* calculated fixation index (F_ST_) estimates. To determine dominant symbiont types, we mapped 2bRAD sequences to a combined symbiont reference composed of transcriptomes from *Symbiodinium* “clades” A and B (Bayer et al. 2012) and “clades” C and D (Ladner, Barshis, and Palumbi 2012) using a custom perl script zooxtype.pl.

Custom scripts are hosted within the 2bRAD GitHub repository (https://github.com/z0on/2bRAD_denovo).

## RESULTS

### 2bRAD Genotyping

An average of 71.9% of reads uniquely mapped to the *M. cavernosa* genome across the 64 2bRAD libraries and an average of 67.3% of sites were covered at >5× sequencing depth. We identified two pairs of clones (7/29 and 17/21; Supp. Fig. 1), which are presumably fragments inadvertently sampled from different parts of the same colony.

**Supplemental Figure1:**
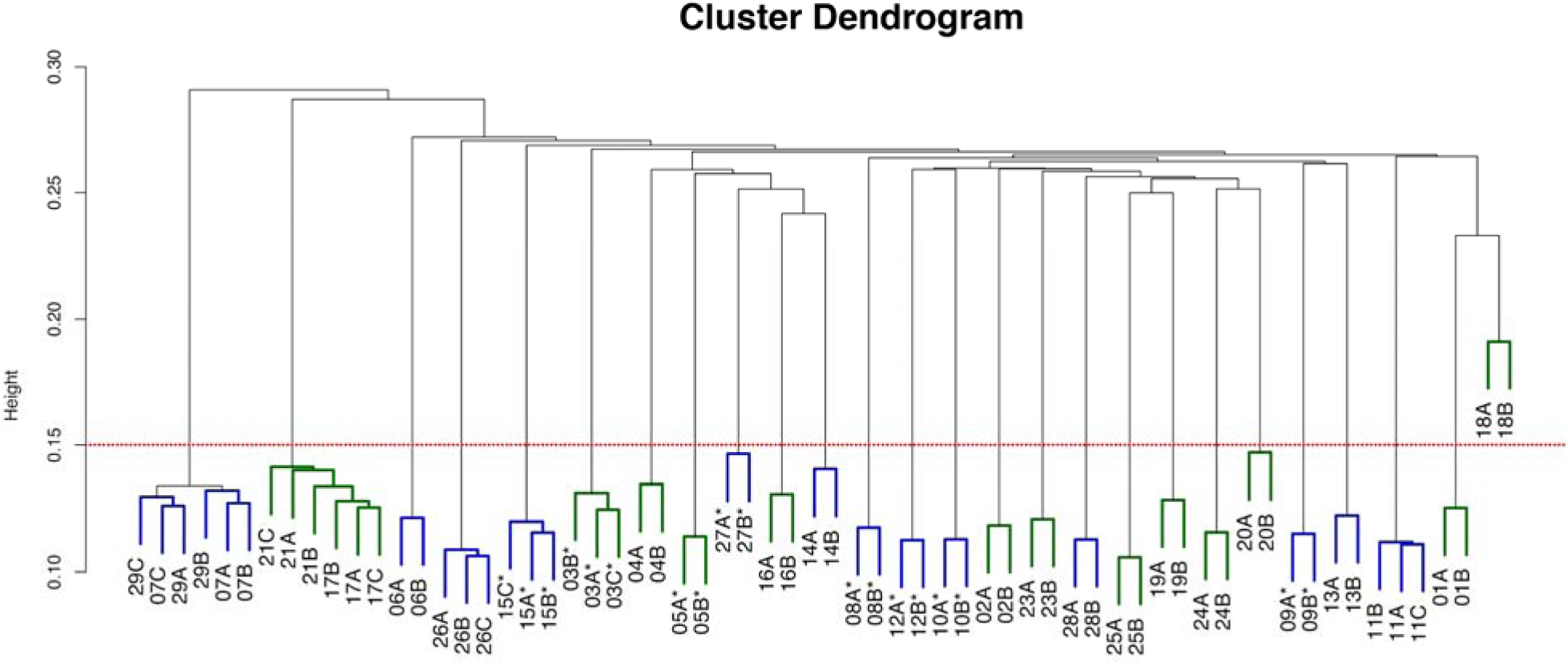
Sample dendrogram based on identity-by-state including clones and genotyping replicates. The sample name includes a number denoting the sample identification at the time of sampling and letter indicating replicate library preparations. Dendrogram color indicates sampling origin: East FGB = blue and West FGB = green. Asterisks indicate *Vibrio* resistance. The dashed red line indicates the threshold for calling distinct genets. Distances <0.15 are presumed clones.

We identified 11,081 SNPs from 26 unique (non-clonal) samples. PCA based on identity-by-state demonstrates a lack of genetic structure between sampling sites or resistance phenotype groups (Supp. Fig. 2).

**Supplemental Figure 2:**
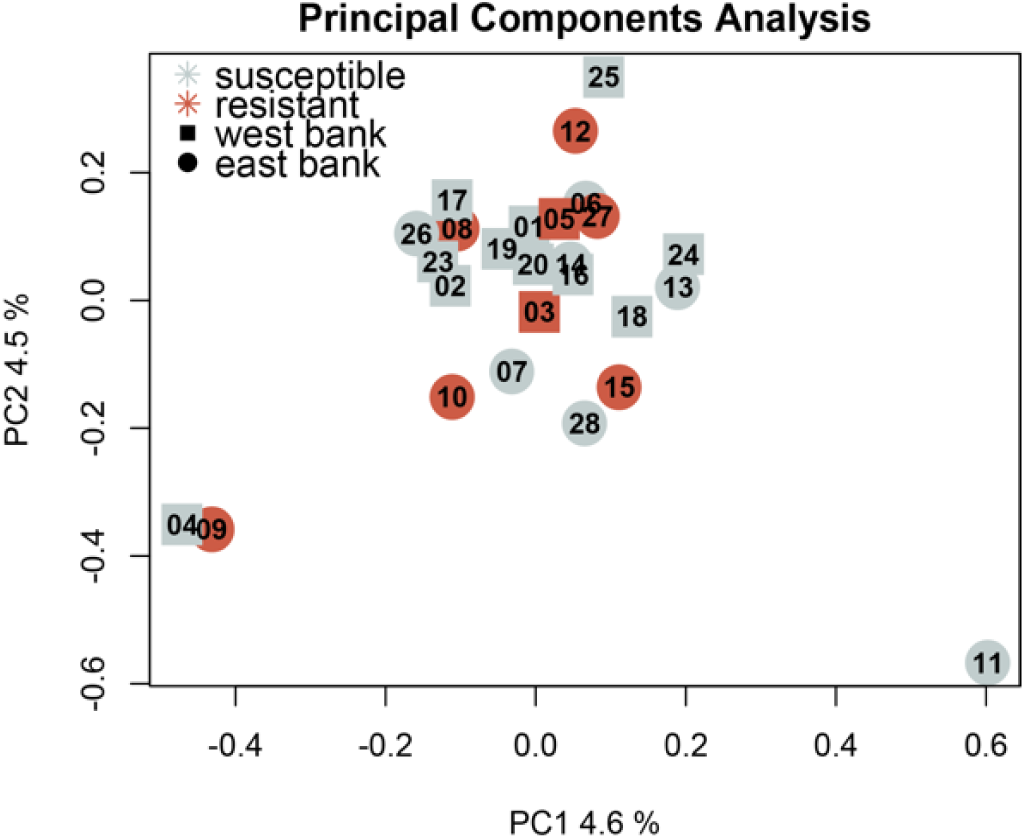
Principal components analysis based on 11,081 genotype probabilities using ANGSD after removing library replicates and clones.

F_ST_ represents the level of genetic differentiation between groups. We calculated weighted F_ST_, which accounts for differences in the numbers of individuals in each group. We observed no genetic differentiation between corals grouped by resistance phenotype (F_ST_ = 0) and little genetic differentiation between East and West FGB origin (F_ST_ = 0.004).

Mapping 2bRAD-seq data to symbiont references determined that all corals were dominated by *Cladocopium* (Supplemental Figure 3).

**Supplemental Figure 3:**
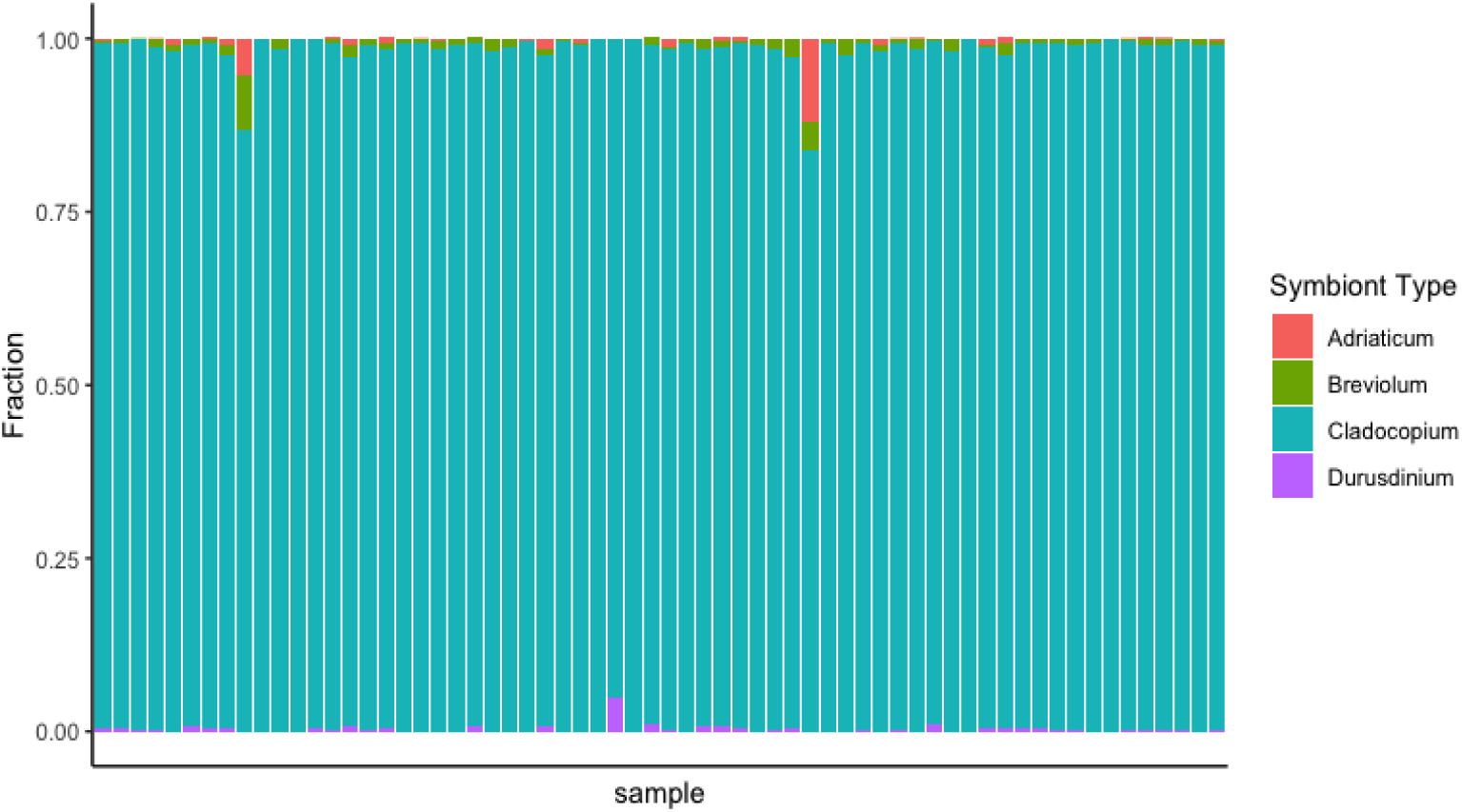
Determination of dominant Symbiodiniaceae types based on 2bRAD-seq data. Columns represent sequencing samples. Colored bars represent the proportion of algal symbiont type according to the legend.

### Bacterial Challenge Survival

Time-of-death was recorded when a coral fragment displayed >50% tissue loss (e.g., Figure 1A). Bacterial challenge significantly increased mortality (p = 2e-16; Figure 1B). Collection site was not associated with differential survival (p = 0.13; Figure 1C).

**Figure 1:**
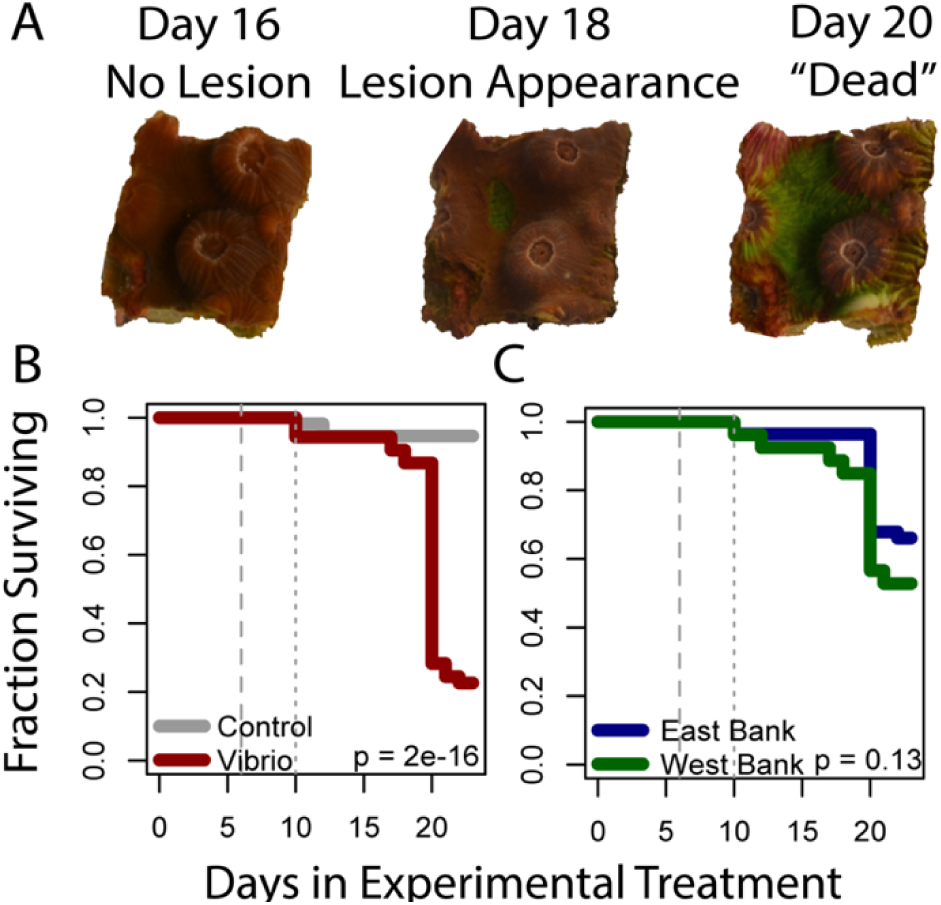
(A) Example lesion progression. (B) Survival of coral fragments in the control (grey) or *Vibrio*-treatment treatments (red). (C) Survival of coral fragments from East FGB (blue) or West FGB (green). P-values correspond to the effect of treatment (B) or collection site (C) in a Cox proportional hazards model. The dashed line at day 6 denotes a shift from 23°C to 29°C. The dotted line at day 10 and a shift from *Vibrio shiloi* to *V. coralliilyticus* exposure.

### Long-term Coral Growth

Across all coral fragments, the average (±SD) long-term calcification rate was 135±61 mg cm^-2^ year^-1^. We observed between 0–30 new polyps on each fragment over the year (mean±SD = 10.9±6.7), which ranged from approximately 0–7 new polyps per cm^2^ of surface area. Neither annual calcification rate nor annual polyp generation was significantly associated with collection site, treatment, or resistance phenotype (Supplemental Figure 4).

**Supplemental Figure 4:**
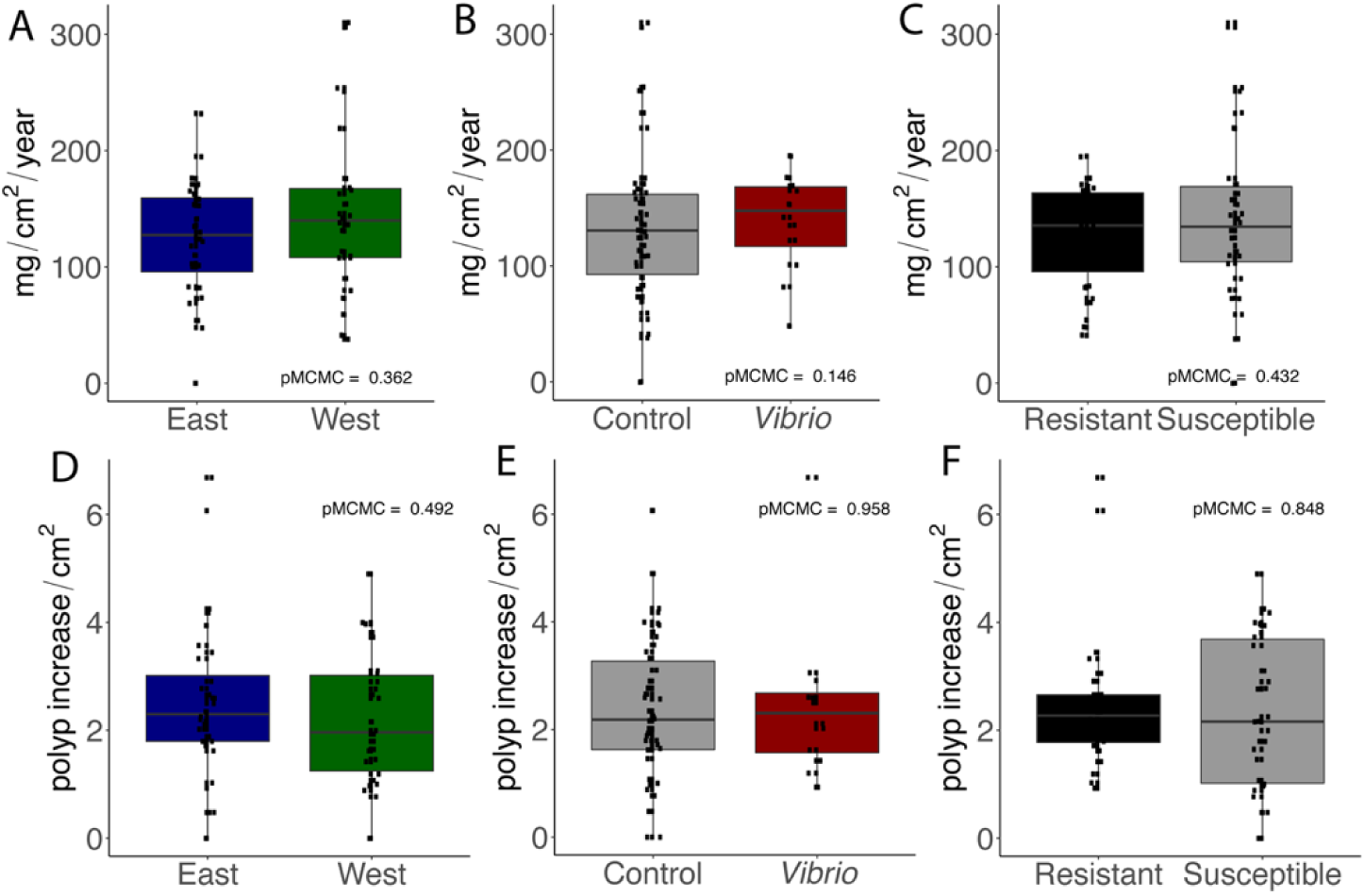
Calcification rate (mg cm^-2^ year^-1^; top) was not associated with collection site (A), treatment (B), or resistance phenotype (C). Polyp generation (new polyps cm^-2^; bottom) was not also associated with collection site (D), treatment (E), or resistance phenotype (F).

### Host Differential Gene Expression

TagSeq yielded an average 319,547 *M. cavernosa* (coral host) gene counts per sample after filtering lowly expressed genes (base mean < 3). The contrast between resistant and susceptible genets yielded only one DEG at FDR = 0.05. This transcript shares substantial homology with a lymphocyte antigen 6H-like gene identified in *Orbicella faveolata* (E-value = 2e-141, identity = 87%) and was more highly expressed in resistant genets (Wald stat = 6.2, FDR = 5.7e-6). No genes were differentially expressed according to sampling origin and only one unannotated gene was significantly associated with annual increase in polyp number (Mcavernosa00313, Wald stat = −4.8, FDR = 0.011).

Seventy DEGs were significantly associated with the rate of calcification in the year following TagSeq library preparation (Supplementary Table 1). Of particular interest, Hairy and Enhancer of Split 1 (Mcavernosa12226, Wald stat = 3.8, FDR = 0.019), Indian hedgehog protein (Mcavernosa00597, Wald stat = 4.1, FDR = 0.016), and A disintegrin and metalloproteinase with thrombospondin motifs 18-like (Mcavernosa26186, Wald stat = 3.6, FDR = 0.045) more highly expressed in fast-growing genets. The top DEGs associated with calcification rate include those encoding a putative vesicular trafficking protein (TOM1-like protein 2) and several uncharacterized proteins (Figure 2).

**Figure 2:**
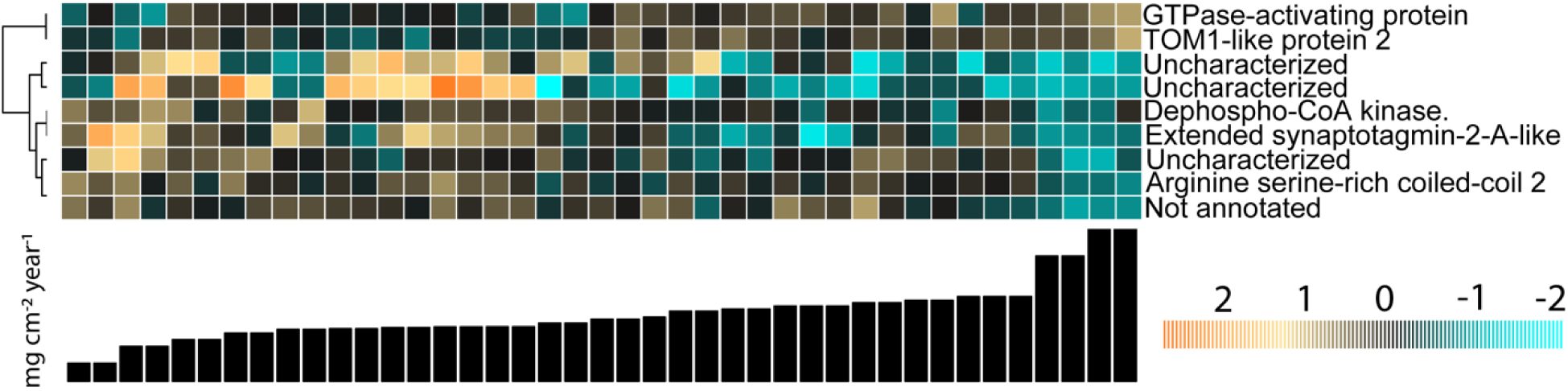
Gene expression associated with annual calcification rate (FDR < 0.01). Heatmap rows are genes and columns are samples. Samples are ordered by calcification rate, as indicated by the bar heights. The color scale is in log^2^-fold change relative to the gene’s mean. Genes are hierarchically clustered based on Pearson’s correlations across samples.

GO enrichment tests yielded 25 biological processes, 22 cellular components, and 5 molecular functions significantly enriched (adjusted *p* < 0.05) with genes associated with calcification rate (Supplemental Fig 5). Among these terms, “cation transport” and “vesicle” were enriched with genes showing higher expression in corals with faster annual calcification rates.

**Supplemental Figure 5:**
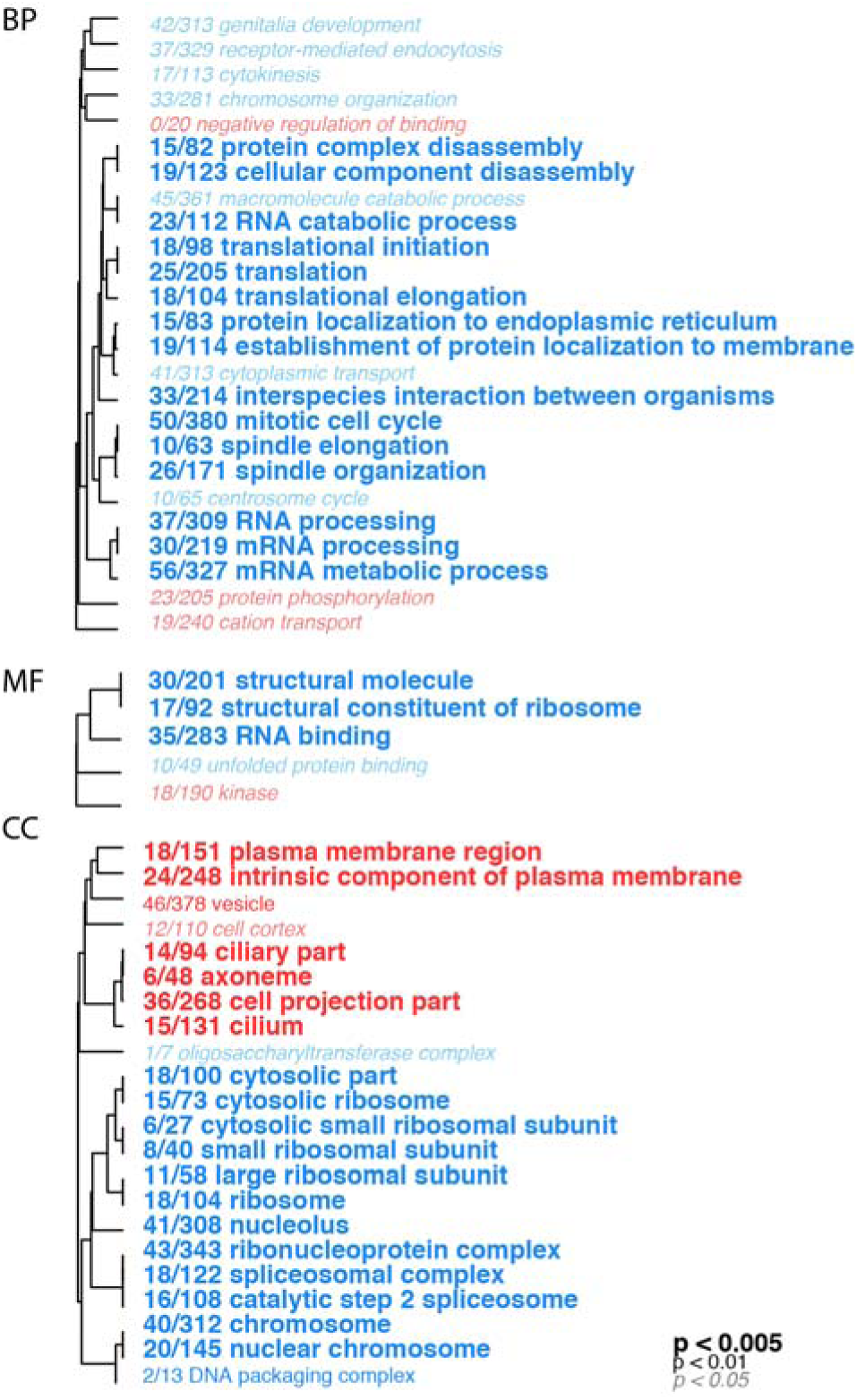
Biological processes (BP), molecular functions (MF), and cellular components (CC) enriched by adjusted p-value generated by testing for association with annual calcification rate. The text color indicates the direction of differential expression according to calcification rate (red = upregulated in faster growing corals; blue = upregulated in slower growing corals). The text size indicates the significance of the term as indicated by the inset key. The fraction preceding the term indicates the number of genes within the term that had an adjusted p-value less than 0.05. Trees indicate gene sharing among gene ontology categories (categories with no branch length between them are subsets of each other).

GO enrichment tests between resistant and susceptible corals yielded 8 biological processes, 20 cellular components, and 5 molecular functions significantly enriched (adjusted *p* < 0.05) among genes associated with disease phenotype. Among these terms, several categories related to cell division (*e.g*., regulation of mitotic cell cycle, DNA integrity checkpoint, kinetochore microtubule) were enriched with genes showing higher expression in resistant corals (Figure 3).

**Figure 3:**
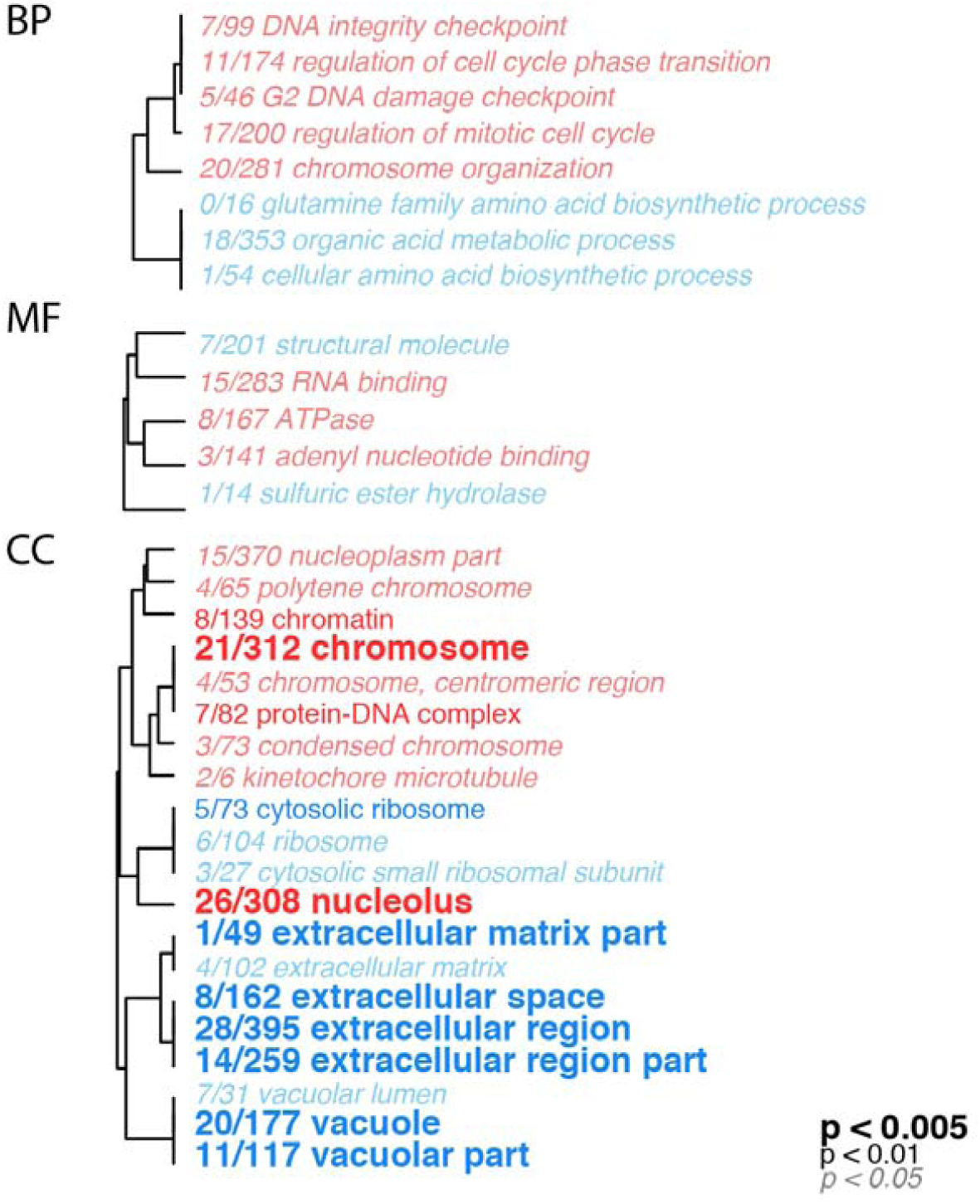
Biological processes (BP), molecular functions (MF), and cellular components (CC) enriched by adjusted p-value generated by testing for association with resistance phenotype. The text color indicates the direction of differential expression between resistant and susceptible genets (red = upregulated in resistant corals; blue = upregulated in susceptible corals). The text size indicates the significance of the term as indicated by the inset key. The fraction preceding the term indicates the number of genes within the term that had an adjusted p-value less than 0.05. Trees indicate gene sharing among gene ontology categories (categories with no branch length between them are subsets of each other).

### Algal Symbiont Gene Expression

TagSeq yielded an average of 9,510 *Cladocopium* (algal symbiont) counts per sample after filtering lowly expressed genes (base mean < 3). No symbiont genes were significantly associated with sampling location, host calcification rate, or host polyp generation (FDR = 0.05). One unannotated gene was significantly associated with host resistance to bacterial challenge (Wald stat = 4.6, FDR = 0.003). GO enrichment analyses did not yield any significantly enriched categories for the symbiont genes.

## DISCUSSION

### No Trade-Offs Between Resistance & Coral Growth

Variation in disease resistance can be explained by differential investment in immunity parameters (Pinzón et al. 2014) that compete for energetic resources with other life-history traits such as growth and reproduction (Leuzinger, Willis, and Anthony 2012). Here we found no association between long-term growth (polyp generation or buoyant weight increase) and disease resistance (Supp. Fig. 4C,F). This result complements previous findings that growth rates in another reef-builder, *Acropora millepora*, were not associated with trade-offs in other health parameters, including survival under *Vibrio* challenge (Wright et al. 2019). We also found that surviving corals demonstrated similar long-term growth rates regardless of whether they received sterile media or *Vibrio* culture during the experimental period (Supp. Fig. 4B,E). These results suggest that growth rates can remain stable after a disease event if a coral can survive and recover from an outbreak. However, back-to-back bleaching events (Head et al. 2019) and multi-year infectious disease outbreaks (Walton, Hayes, and Gilliam 2018) limit the amount of a time a coral can recover before the next life-threatening challenge.

### Genomic Associations with Disease Resistance

Gene expression analysis revealed subtle differences in pre-exposure transcriptomic states between corals that subsequently demonstrated resistance or susceptibility to *Vibrio* challenge. Only one transcript, which shares extensive homology with a lymphocyte antigen-6 (Ly6) gene, passed the genome-wide significance threshold of FDR=0.05; it was upregulated in resistant corals prior to bacterial challenge (Supplementary Data 1). Genes belonging to the Ly6 family play various roles across metazoans, such as epithelial barrier formation in *Drosophila* (Hijazi et al. 2011) and neutrophil migration in mammals (Lee et al. 2013). In mouse epithelial cells, expression of a Ly6 protein (Lypd8) promotes gut homeostasis and prevents pathogen attachment (Okumura et al. 2020). Given that many coral diseases are associated with loss of tissue structure and bacterial infiltration (Ainsworth et al. 2007), future studies should explore the potential role of this gene family in promoting coral tissue integrity upon bacterial challenge.

Mitotic activity including spindle formation and cell cycle phase regulation were enriched among upregulated genes in resistant individuals (Fig. 3), possibly indicating an abundance of a specific population of proliferative cells or higher cell division rates in resistant individuals. To test these hypotheses, single-cell transcriptomics may reveal differences in activated cell populations between resistant and susceptible genets. Alternatively, senescence may explain differences in cell growth rates (Rinkevich and Loya 1986). Colony age can explain differences in disease susceptibility, as has been reported in *Acropora palmata* affected with White-Pox disease (Muller and van Woesik 2014). All coral cells of a clonal genet have the same age since sexual recruitment (i.e., chronological age), but soft tissues across the colony have likely experienced different numbers of cell divisions (i.e., replicative age). Across colonies of the massive reef-builder *Porites* with an average age of 41 years, the average polyp age was only 2–3 years (Darke and Barnes 1993). Future investigations of the impacts of aging and age-related cell turnover rates (Petralia, Mattson, and Yao 2014) on disease susceptibility in corals should evaluate markers of replicative age, such as telomere length or somatic mutation accumulation (Barfield, Aglyamova, and Matz 2016).

The small sample size of this study precludes investigation of the genomic architecture underlying disease resistance, though *in situ* disease transmission experiments provide evidence for a genetic basis to disease resistance in some species of reef-building corals (Vollmer and Kline 2008; Libro and Vollmer 2016). A recent study that identified dozens of genetic variants associated with resistance to *Vibrio* infection in a flatfish used phenotype data from thousands of fish and whole-genome resequencing for over 500 individuals (Zhou et al. 2019). Conducting experiments at this scale in threatened coral species presents considerable challenges, though genomic predictors for thermal tolerance in corals have been possible through low-coverage sequencing from minimally invasive tissue samples from hundreds of adult colonies *in situ* (Fuller et al. 2020) and genome-wide SNP analysis of coral larvae produced through sexual reproduction from experimentally selected parent colonies (Quigley, Bay, and van Oppen 2020). Our estimates of F_ST_ between corals from the East and West FGB match previous studies (Studivan and Voss 2018) and support models of high gene flow through larval dispersal in the region (S. W. Davies et al. 2017; Goodbody-Gringley, Woollacott, and Giribet 2012).

### Predictive Gene Expression Associated with Long-Term Calcification

Transcripts homologous to Hairy and Enhancer of Split (HES), Indian hedgehog protein (IHH), a disintegrin and metalloproteinase with thrombospondin motifs (ADAMTS) and were more highly expressed in corals with faster growth rates (Supp. Data 1). HES and IHH regulate bone mass in mammals (Zanotti, Smerdel-Ramoya, and Canalis 2011; Deng et al. 2018), but their role in coral biology is currently unclear. ADAMTS enzymes have diverse roles in extracellular matrix remodeling and tissue morphogenesis (reviewed in Kelwick et al. 2015). Two ADAMTS transcripts were also upregulated in *Stylophora pistillata* treated with high calcium concentrations that subsequently increased calcification rates (Gutner-Hoch et al. 2017). A study examining gene expression patterns predicting growth in *Acropora hyacinthus* transplanted to new environments also identified an ADAMTS transcript among genes upregulated in faster growing corals (Bay and Palumbi 2017). Other similarities in predictive transcriptomic results between this study and Bay & Palumbi (2017) include the upregulation of TOM1-like protein 2 and lectin in fast-growing corals. Given their association with long-term growth across multiple coral species, reefs, and sampling timepoints (Seneca and Palumbi 2015), these genes represent prime candidates for validation as potential predictive growth biomarkers.

### Conclusions

We demonstrate intraspecific variation in pathogen resistance in a reef-building coral from an isolated marine sanctuary with no documented instance of coral disease. Understanding the immediate and long-term consequences of bacterial pathogen exposure is especially important given the potential impact of this sanctuary as a larval source to restore disease-degraded Caribbean coral populations. The presence of resistant genets and lack of trade-offs between resistance and growth under these laboratory conditions provide hope that this coral population may be able to withstand some bacterial challenge. However, ever-worsening ocean conditions threaten marine organisms with multiple concurrent stressors. The health of coral reefs ultimately relies on global action to mitigate the effects of climate change.

## ACKNOWLEDGEMENTS

The authors are grateful for the NOAA research team at FGBNMS, including Emma Hickerson and the crew of the R/V Manta for field assistance. These corals were collected under permit FGBNMS-2014-0013. We thank the Texas Advanced Computing Center for computational resources and to Dr. Laura Katz for mentorship.

## DATA ACCESSIBILITY

Sequences are deposited in the SRA PRJNA355872 (TagSeq) and PRJNA655196 (2bRAD). Phenotype data and analysis scripts are hosted at https://github.com/rachelwright8/McavSusceptibility. Custom 2bRAD scripts are hosted within the 2bRAD GitHub repository (https://github.com/z0on/2bRAD_denovo).

## AUTHORS’ CONTRIBUTIONS

ERK analyzed the phenotypic and gene expression data and contributed to manuscript preparation. RSS participated in data analysis and contributed to manuscript preparation. MVM coordinated the study and critically revised the manuscript. RMW conceived of and designed the study, collected coral specimens, carried out the bacterial challenge and molecular work, analyzed the 2bRAD sequence data, and drafted the manuscript.

## REFERENCES

Ainsworth, T. D., E. Kramasky-Winter, Y. Loya, O. Hoegh-Guldberg, and M. Fine. 2007. “Coral Disease Diagnostics: What’s between a Plague and a Band?” Applied and Environmental Microbiology 73 (3): 981–92.

Aronson, R. B., W. F. Precht, and I. G. Macintyre. 1998. “Extrinsic Control of Species Replacement on a Holocene Reef in Belize: The Role of Coral Disease.” Coral Reefs. https://doi.org/10.1007/s003380050122.

Barfield, Sarah, Galina V. Aglyamova, and Mikhail V. Matz. 2016. “Evolutionary Origins of Germline Segregation in Metazoa: Evidence for a Germ Stem Cell Lineage in the Coral Orbicella Faveolata (Cnidaria, Anthozoa).” Proceedings. Biological Sciences / The Royal Society 283 (1822). https://doi.org/10.1098/rspb.2015.2128.

Bayer, Till, Manuel Aranda, Shinichi Sunagawa, Lauren K. Yum, Michael K. Desalvo, Erika Lindquist, Mary Alice Coffroth, Christian R. Voolstra, and Mónica Medina. 2012. “Symbiodinium Transcriptomes: Genome Insights into the Dinoflagellate Symbionts of Reef-Building Corals.” PloS One 7 (4): e35269.

Bay, Rachael A., and Stephen R. Palumbi. 2017. “Transcriptome Predictors of Coral Survival and Growth in a Highly Variable Environment.” Ecology and Evolution 7 (13): 4794–4803.

Benjamini, Yoav, and Yosef Hochberg. 1995. “Controlling the False Discovery Rate: A Practical and Powerful Approach to Multiple Testing.” Journal of the Royal Statistical Society: Series B (Methodological). https://doi.org/10.1111/j.2517-6161.1995.tb02031.x.

Chimetto Tonon, Luciane A., Janelle R. Thompson, Ana P. B. Moreira, Gizele D. Garcia, Kevin Penn, Rachelle Lim, Roberto G. S. Berlinck, Cristiane C. Thompson, and Fabiano L. Thompson. 2017. “Quantitative Detection of Active Vibrios Associated with White Plague Disease in Corals.” Frontiers in Microbiology 8 (November): 2272.

C., Jorge H. Pinzón, Jorge H. Pinzón C., Joshuah Beach-Letendre, Ernesto Weil, and Laura D. Mydlarz. 2014. “Relationship between Phylogeny and Immunity Suggests Older Caribbean Coral Lineages Are More Resistant to Disease.” PLoS ONE. https://doi.org/10.1371/journal.pone.0104787.

Darke, W. M., and D. J. Barnes. 1993. “Growth Trajectories of Corallites and Ages of Polyps in Massive Colonies of Reef-Building Corals of the Genus Porites.” Marine Biology. https://doi.org/10.1007/bf00345677.

Davies, P. Spencer, and P. Spencer Davies. 1989. “Short-Term Growth Measurements of Corals Using an Accurate Buoyant Weighing Technique.” Marine Biology. https://doi.org/10.1007/bf00428135.

Davies, Sarah W., Justin B. Ries, Adrian Marchetti, and Karl D. Castillo. 2018. “Symbiodinium Functional Diversity in the Coral Siderastrea Siderea Is Influenced by Thermal Stress and Reef Environment, but Not Ocean Acidification.” Frontiers in Marine Science 5: 150.

Davies, Sarah W., Marie E. Strader, Johnathan T. Kool, Carly D. Kenkel, and Mikhail V. Matz. 2017. “Modeled Differences of Coral Life-History Traits Influence the Refugium Potential of a Remote Caribbean Reef.” Coral Reefs. https://doi.org/10.1007/s00338-017-1583-8.

Deng, Ang, Hongqi Zhang, Minyu Hu, Shaohua Liu, Yuxiang Wang, Qile Gao, and Chaofeng Guo. 2018. “The Inhibitory Roles of Ihh Downregulation on Chondrocyte Growth and Differentiation.” Experimental and Therapeutic Medicine 15 (1): 789–94.

Fuess, Lauren E., Jorge H. Pinzón C, Ernesto Weil, Robert D. Grinshpon, and Laura D. Mydlarz. 2017. “Life or Death: Disease-Tolerant Coral Species Activate Autophagy Following Immune Challenge.” Proceedings. Biological Sciences / The Royal Society 284 (1856). https://doi.org/10.1098/rspb.2017.0771.

Fuller, Zachary L., Veronique J. L. Mocellin, Luke A. Morris, Neal Cantin, Jihanne Shepherd, Luke Sarre, Julie Peng, et al. 2020. “Population Genetics of the Coral : Toward Genomic Prediction of Bleaching.” Science 369 (6501). https://doi.org/10.1126/science.aba4674.

Fuller, Z., Yi, L., and Matz, M. n.d. “The Genome of Montastrea Cavernosa. Available from https://matzlab.weebly.com/data--Code.html.”

Goodbody-Gringley, Gretchen, Robert M. Woollacott, and Gonzalo Giribet. 2012. “Population Structure and Connectivity in the Atlantic Scleractinian Coral Montastraea Cavernosa (Linnaeus, 1767).” Marine Ecology 33 (1): 32–48.

Gordon, Assaf, G. J. Hannon, and Others. 2010. “Fastx-Toolkit.” FASTQ/A Short-Reads Preprocessing Tools (unpublished) http://hannonlab.Cshl.Edu/fastx_toolkit5.

Gutner-Hoch, Eldad, Hiba Waldman Ben-Asher, Ruth Yam, Aldo Shemesh, and Oren Levy. 2017. “Identifying Genes and Regulatory Pathways Associated with the Scleractinian Coral Calcification Process.” PeerJ 5 (July): e3590.

Hadfield, Jarrod D. 2010. “MCMC Methods for Multi-Response Generalized Linear Mixed Models: The MCMCglmm R Package.” Journal of Statistical Software 33 (2): 1–22.

Head, Catherine E. I., Daniel T. I. Bayley, Gwilym Rowlands, Ronan C. Roche, David M. Tickler, Alex D. Rogers, Heather Koldewey, John R. Turner, and Dominic A. Andradi-Brown. 2019. “Coral Bleaching Impacts from Back-to-Back 2015–2016 Thermal Anomalies in the Remote Central Indian Ocean.” Coral Reefs 38 (4): 605–18.

Hijazi, Assia, Marc Haenlin, Lucas Waltzer, and Fernando Roch. 2011. “The Ly6 Protein Coiled Is Required for Septate Junction and Blood Brain Barrier Organisation in Drosophila.” PLoS ONE. https://doi.org/10.1371/journal.pone.0017763.

Hobbs, Jean-Paul A., Ashley J. Frisch, Stephen J. Newman, and Corey B. Wakefield. 2015. “Selective Impact of Disease on Coral Communities: Outbreak of White Syndrome Causes Significant Total Mortality of Acropora Plate Corals.” PloS One 10 (7): e0132528.

Jin, Young K., Petra Lundgren, Adrian Lutz, Jean-Baptiste Raina, Emily J. Howells, Allison S. Paley, Bette L. Willis, and Madeleine J. H. van Oppen. 2016. “Genetic Markers for Antioxidant Capacity in a Reef-Building Coral.” Science Advances 2 (5): e1500842.

Johnston, Michelle A., John A. Embesi, Ryan J. Eckert, Marissa F. Nuttall, Emma L. Hickerson, and George P. Schmahl. 2016. “Persistence of Coral Assemblages at East and West Flower Garden Banks, Gulf of Mexico.” Coral Reefs. https://doi.org/10.1007/s00338-016-1452-x.

Johnston, Michelle A., Marissa F. Nuttall, Ryan J. Eckert, Raven D. Blakeway, Travis K. Sterne, Emma L. Hickerson, George P. Schmahl, Michael T. Lee, James MacMillan, and John A. Embesi. 2019. “Localized Coral Reef Mortality Event at East Flower Garden Bank, Gulf of Mexico.” Bulletin of Marine Science. https://doi.org/10.5343/bms.2018.0057.

Kauffmann, Audrey, Robert Gentleman, and Wolfgang Huber. 2008. “arrayQualityMetrics—a Bioconductor Package for Quality Assessment of Microarray Data.” Bioinformatics 25 (3): 415–16.

Kealoha, Andrea K., Shawn M. Doyle, Kathryn E. F. Shamberger, Jason B. Sylvan, Robert D. Hetland, and Steven F. DiMarco. 2020. “Localized Hypoxia May Have Caused Coral Reef Mortality at the Flower Garden Banks.” Coral Reefs. https://doi.org/10.1007/s00338-019-01883-9.

Kelwick, Richard, Ines Desanlis, Grant N. Wheeler, and Dylan R. Edwards. 2015. “The ADAMTS (A Disintegrin and Metalloproteinase with Thrombospondin Motifs) Family.” Genome Biology 16 (May): 113.

Kolde, Raivo. 2012. “Pheatmap: Pretty Heatmaps.” R Package Version 1 (2).

Korneliussen, Thorfinn Sand, Anders Albrechtsen, and Rasmus Nielsen. 2014. “ANGSD: Analysis of Next Generation Sequencing Data.” BMC Bioinformatics 15 (November): 356.

Ladner, Jason T., Daniel J. Barshis, and Stephen R. Palumbi. 2012. “Protein Evolution in Two Co-Occurring Types of Symbiodinium: An Exploration into the Genetic Basis of Thermal Tolerance in Symbiodiniumclade D.” BMC Evolutionary Biology 12 (1): 217.

Langmead, B., and S. L. Salzberg. 2013. “Langmead. 2013. Bowtie2.” Nature Methods 9: 357–59.

Lee, Pui Y., Jun-Xia Wang, Emilio Parisini, Christopher C. Dascher, and Peter A. Nigrovic. 2013. “Ly6 Family Proteins in Neutrophil Biology.” Journal of Leukocyte Biology 94 (4): 585–94.

Leuzinger, Sebastian, Bette L. Willis, and Kenneth R. N. Anthony. 2012. “Energy Allocation in a Reef Coral under Varying Resource Availability.” Marine Biology 159 (1): 177–86.

Libro, Silvia, and Steven V. Vollmer. 2016. “Genetic Signature of Resistance to White Band Disease in the Caribbean Staghorn Coral Acropora Cervicornis.” PloS One 11 (1): e0146636.

Love, Michael I., Wolfgang Huber, and Simon Anders. 2014. “Moderated Estimation of Fold Change and Dispersion for RNA-Seq Data with DESeq2.” Genome Biology 15 (12): 550.

Manzello, Derek P., Ian C. Enochs, Graham Kolodziej, and Renée Carlton. 2015. “Coral Growth Patterns of Montastraea Cavernosa and Porites Astreoides in the Florida Keys: The Importance of Thermal Stress and Inimical Waters.” Journal of Experimental Marine Biology and Ecology. https://doi.org/10.1016/j.jembe.2015.06.010.

Meyer, E., G. V. Aglyamova, and M. V. Matz. 2011. “Profiling Gene Expression Responses of Coral Larvae (Acropora Millepora) to Elevated Temperature and Settlement Inducers Using a Novel RNA-Seq Procedure.” Molecular Ecology 20 (17): 3599–3616.

Muller, Erinn M., and Robert van Woesik. 2014. “Genetic Susceptibility, Colony Size, and Water Temperature Drive White-Pox Disease on the Coral Acropora Palmata.” PloS One 9 (11): e110759.

Okumura, Ryu, Toshio Kodama, Chiao-Ching Hsu, Benjamin Heller Sahlgren, Shota Hamano, Takashi Kurakawa, Tetsuya Iida, and Kiyoshi Takeda. 2020. “Lypd8 Inhibits Attachment of Pathogenic Bacteria to Colonic Epithelia.” Mucosal Immunology 13 (1): 75–85.

Oppen, Madeleine J. H. van, Ruth D. Gates, Linda L. Blackall, Neal Cantin, Leela J. Chakravarti, Wing Y. Chan, Craig Cormick, et al. 2017. “Shifting Paradigms in Restoration of the World’s Coral Reefs.” Global Change Biology 23 (9): 3437–48.

Petralia, Ronald S., Mark P. Mattson, and Pamela J. Yao. 2014. “Aging and Longevity in the Simplest Animals and the Quest for Immortality.” Ageing Research Reviews. https://doi.org/10.1016/j.arr.2014.05.003.

Precht, William F., Brooke E. Gintert, Martha L. Robbart, Ryan Fura, and Robert van Woesik. 2016. “Unprecedented Disease-Related Coral Mortality in Southeastern Florida.” Scientific Reports 6 (August): 31374.

Quigley, Kate M., Line K. Bay, and Madeleine J. H. van Oppen. 2020. “Genome-Wide SNP Analysis Reveals an Increase in Adaptive Genetic Variation through Selective Breeding of Coral.” Molecular Ecology 29 (12): 2176–88.

Rinkevich, B., and Y. Loya. 1986. “Senescence and Dying Signals in a Reef Building Coral.” Experientia 42 (3): 320–22.

Rippe, John P., Nicola G. Kriefall, Sarah W. Davies, and Karl D. Castillo. 2019. “Differential Disease Incidence and Mortality of Inner and Outer Reef Corals of the Upper Florida Keys in Association with a White Syndrome Outbreak.” Bulletin of Marine Science. https://doi.org/10.5343/bms.2018.0034.

Rose, Noah H., Francois O. Seneca, and Stephen R. Palumbi. 2016. “Gene Networks in the Wild: Identifying Transcriptional Modules That Mediate Coral Resistance to Experimental Heat Stress.” Genome Biology and Evolution. https://doi.org/10.1093/gbe/evv258.

Schneider, Caroline A., Wayne S. Rasband, and Kevin W. Eliceiri. 2012. “NIH Image to ImageJ: 25 Years of Image Analysis.” Nature Methods 9 (7): 671–75.

Seneca, Francois O., and Stephen R. Palumbi. 2015. “The Role of Transcriptome Resilience in Resistance of Corals to Bleaching.” Molecular Ecology. https://doi.org/10.1111/mec.13125.

Studivan, M. S., and J. D. Voss. 2018. “Population Connectivity among Shallow and Mesophotic Montastraea Cavernosa Corals in the Gulf of Mexico Identifies Potential for Refugia.” Coral Reefs 37 (4): 1183–96.

Team, R Core. 2019. “A Language and Environment for Statistical Computing. Vienna, Austria: R Foundation for Statistical Computing.” URL https://www.R-Project.Org.

Vollmer, Steven V., and David I. Kline. 2008. “Natural Disease Resistance in Threatened Staghorn Corals.” PloS One 3 (11): e3718.

Walton, Charles J., Nicole K. Hayes, and David S. Gilliam. 2018. “Impacts of a Regional, Multi-Year, Multi-Species Coral Disease Outbreak in Southeast Florida.” Frontiers in Marine Science. https://doi.org/10.3389/fmars.2018.00323.

Wang, Shi, Eli Meyer, John K. McKay, and Mikhail V. Matz. 2012. “2b-RAD: A Simple and Flexible Method for Genome-Wide Genotyping.” Nature Methods 9 (8): 808–10.

Weil, Ernesto, Garriet Smith, and Diego L. Gil-Agudelo. 2006. “Status and Progress in Coral Reef Disease Research.” Diseases of Aquatic Organisms 69 (1): 1–7.

Williams, D. E., and M. W. Miller. 2012. “Attributing Mortality among Drivers of Population Decline in Acropora Palmata in the Florida Keys (USA).” Coral Reefs. https://doi.org/10.1007/s00338-011-0847-y.

Wright, Rachel M., Galina V. Aglyamova, Eli Meyer, and Mikhail V. Matz. 2015. “Gene Expression Associated with White Syndromes in a Reef Building Coral, Acropora Hyacinthus.” BMC Genomics 16 (May): 371.

Wright, Rachel M., Carly D. Kenkel, Carly E. Dunn, Erin N. Shilling, Line K. Bay, and Mikhail V. Matz. 2017. “Intraspecific Differences in Molecular Stress Responses and Coral Pathobiome Contribute to Mortality under Bacterial Challenge in Acropora Millepora.” Scientific Reports. https://doi.org/10.1038/s41598-017-02685-1.

Wright, Rachel M., Hanaka Mera, Carly D. Kenkel, Maria Nayfa, Line K. Bay, and Mikhail V. Matz. 2019. “Positive Genetic Associations among Fitness Traits Support Evolvability of a Reef-Building Coral under Multiple Stressors.” Global Change Biology 25 (10): 3294–3304.

Zanotti, Stefano, Anna Smerdel-Ramoya, and Ernesto Canalis. 2011. “HES1 (hairy and Enhancer of Split 1) Is a Determinant of Bone Mass.” The Journal of Biological Chemistry 286 (4): 2648–57.

Zhou, Qian, Zhencheng Su, Yangzhen Li, Yang Liu, Lei Wang, Sheng Lu, Shuanyan Wang, et al. 2019. “Genome-Wide Association Mapping and Gene Expression Analyses Reveal Genetic Mechanisms of Disease Resistance Variations in Cynoglossus Semilaevis.” Frontiers in Genetics 10 (November): 1167.

