## Supplementary figures and images for "Gene expression associated with disease resistance and long-term growth in a reef-building coral"

### Supplemental Figure 1

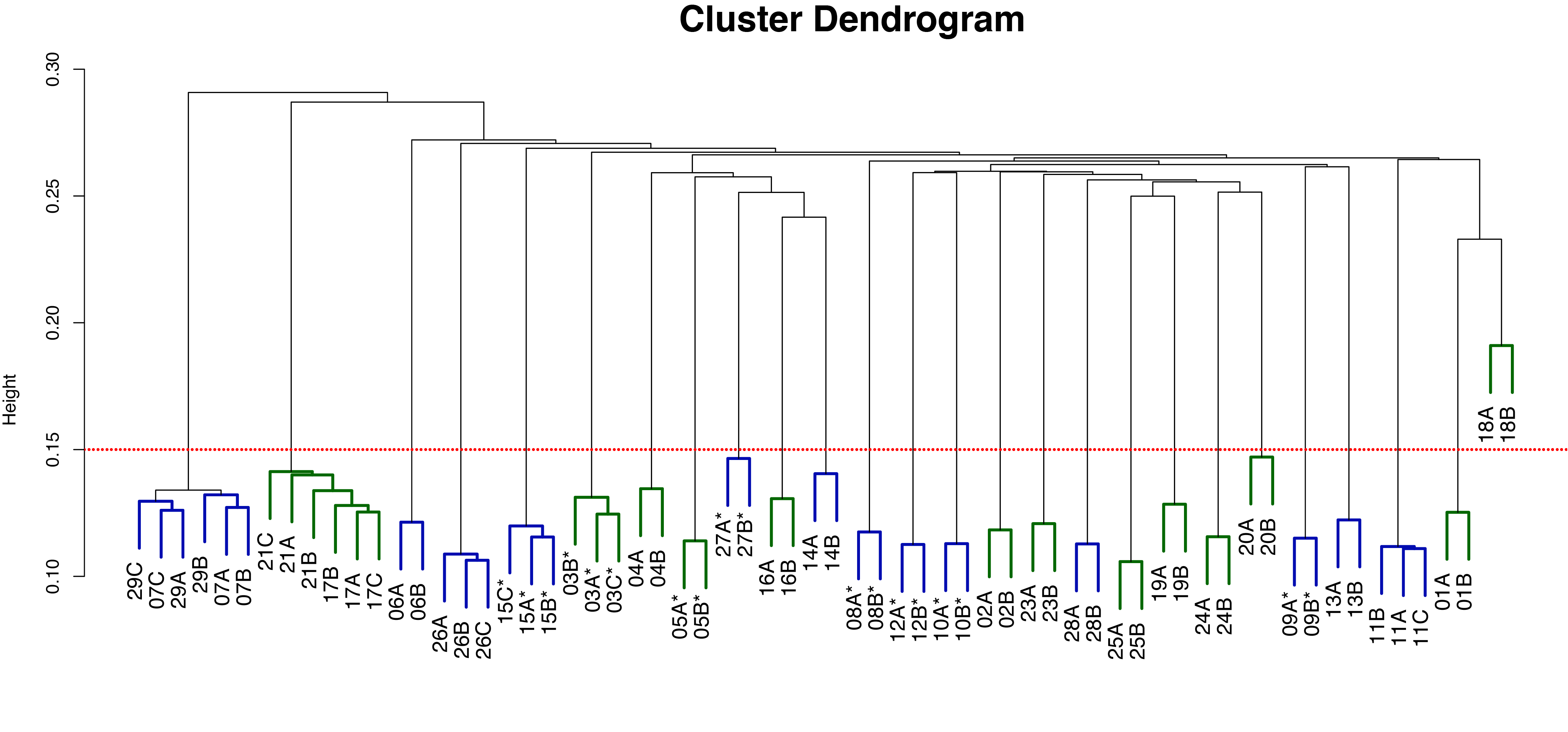

### Supplemental Figure 2

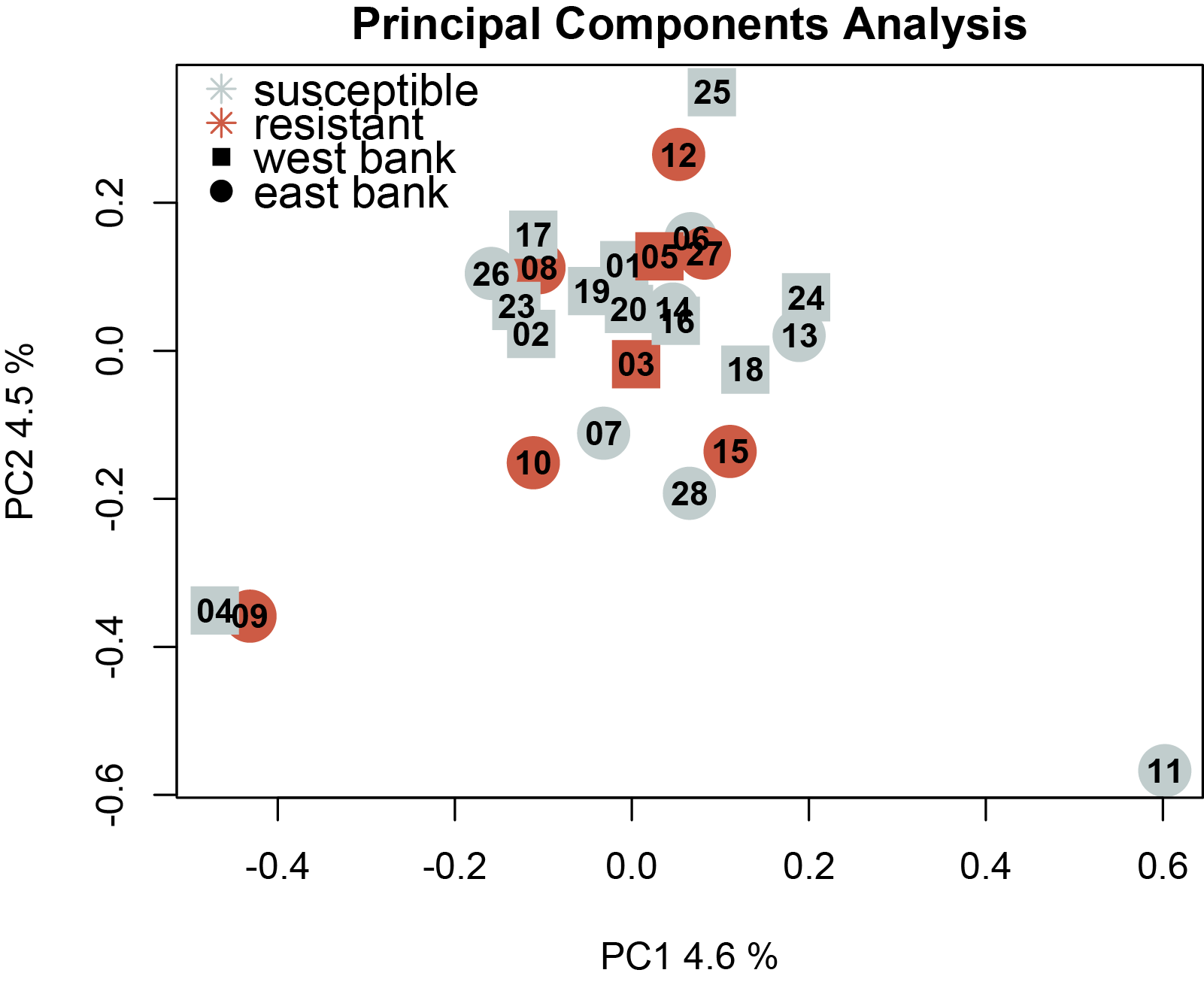

### Supplemental Figure 3

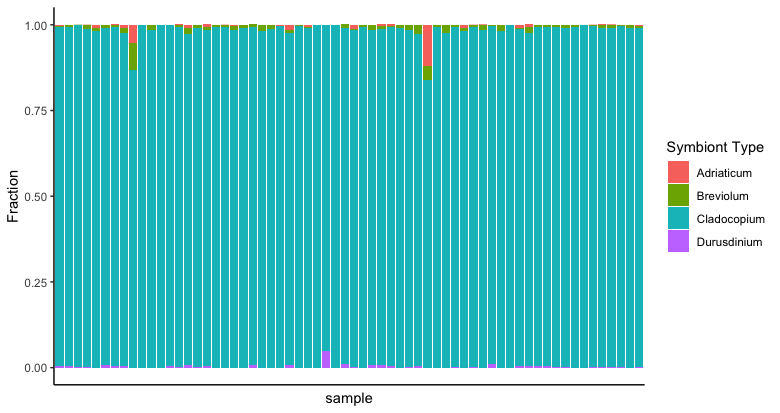

### Supplemental Figure 4

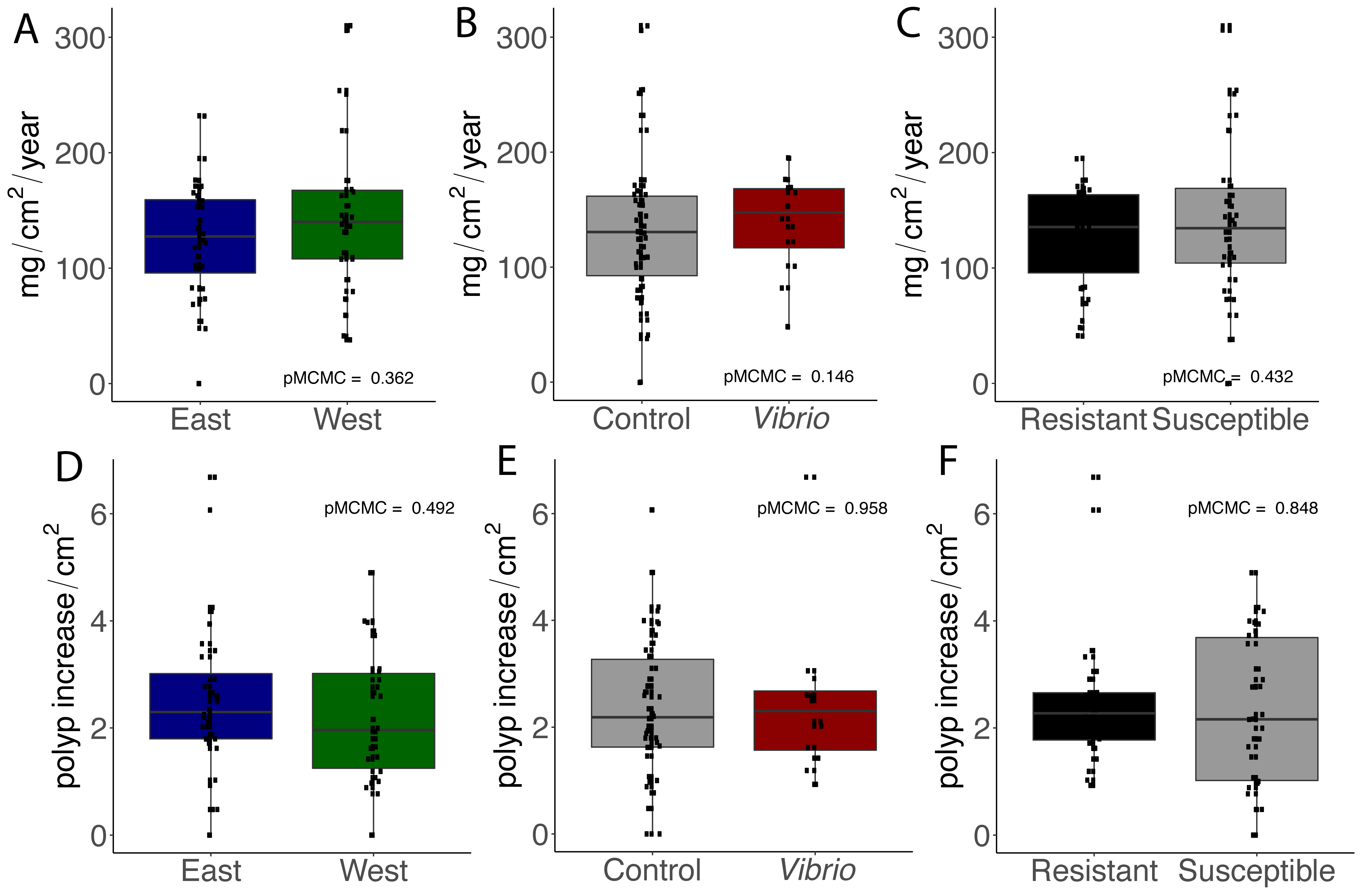

### Supplemental Figure 5

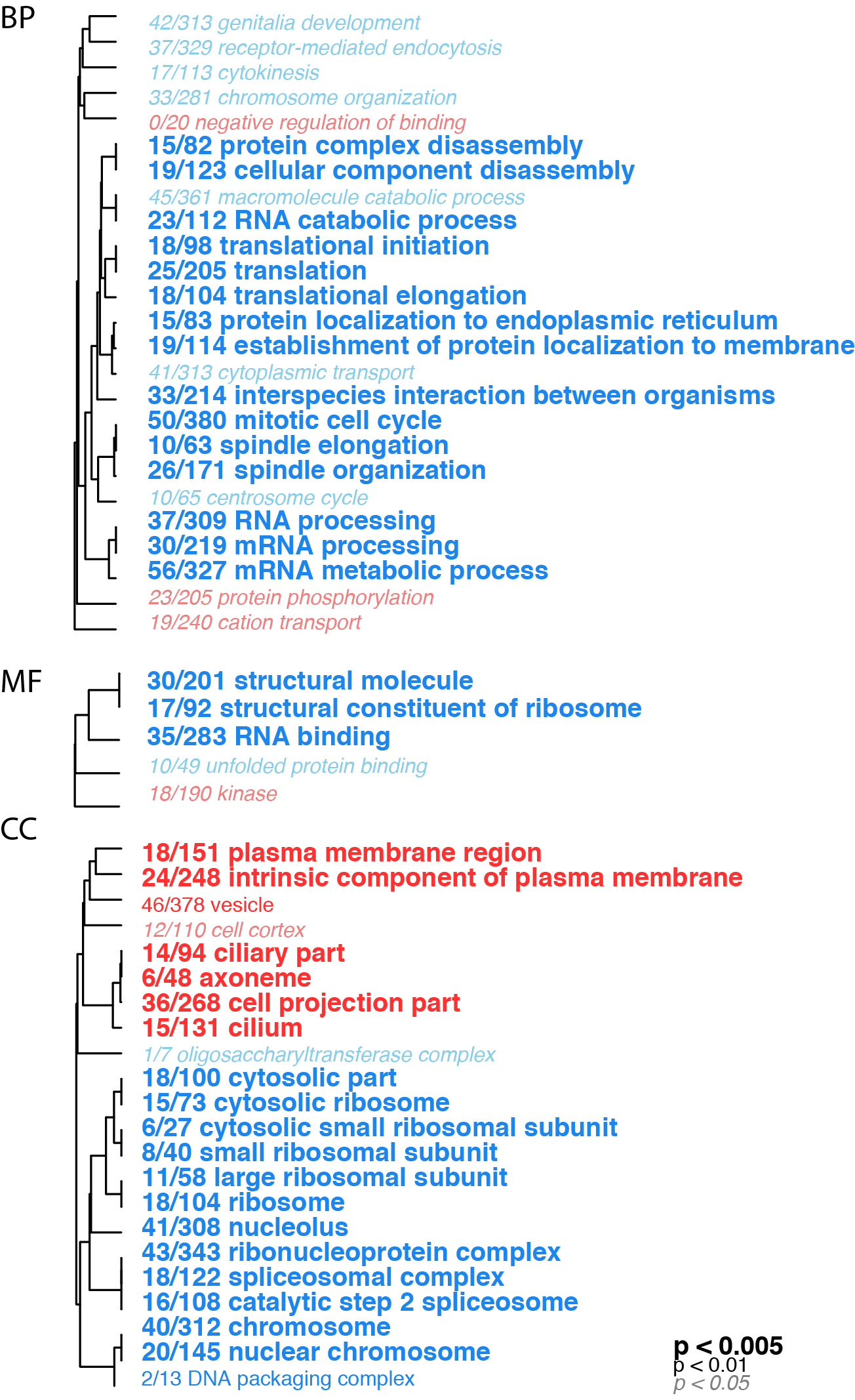
